# Dosage compensation and sexual conflict in female heterogametic methylomes

**DOI:** 10.1101/2024.10.02.616244

**Authors:** Marianthi Tangili, Joanna Sudyka, Fabricio Furni, Per J. Palsbøll, Simon Verhulst

**Affiliations:** Groningen Institute for Evolutionary Life Sciences, University of Groningen, Groningen, The Netherlands; Institute of Environmental Sciences, Jagiellonian University, Kraków, Poland; Center for Coastal Studies, Provincetown, Massachusetts, USA

## Abstract

DNA methylation (DNAm) suppresses gene expression and contributes to dosage compensation in mammals but whether DNAm plays a similar role in female ZW chromosome heterogametic species remains unresolved. We assessed chromosome-level DNAm using whole genome bisulphite sequencing in two avian species, zebra finches and jackdaws. Dosage compensation by DNAm would result in higher and more variable DNAm level in males relative to females on the Z chromosome. However, we found that the level of DNAm and its variance on the Z chromosome was *lower* in males. Moreover, male Z chromosome-based gene promoters were more frequently hypomethylated compared to females, indicating absence of upregulation on a gene-by-gene basis across the female Z chromosome. We suggest our findings reveal mitigation of an intra-genomic sexual conflict, with females suppressing expression of Z chromosome-based genes that benefit male but not female fitness. W was the most methylated chromosome, but hypermethylation on the W chromosome was mostly confined to intergenic regions, presumably resulting in the downregulation of transposable elements known to comprise a large part of the W chromosome. Thus, DNAm is involved in the development of sex-dependent phenotypes, but dosage compensation is achieved through other mechanisms.

## Introduction

Sex-chromosome differentiation drives the development and evolution of sexually dimorphic phenotypes in species with chromosomal sex determination^1^. The Y and W sex chromosomes in species with XY or ZW sex-determination systems are degenerated^2^, containing few genes. This results in a gene imbalance, as the heterogametic sex is homozygous for genes solely located on the X or Z chromosome, whereas the homogametic sex has two copies of one of the sex chromsomes^3,4^. Dosage compensation is a mechanism that addresses potential sex-chromosome gene imbalance between the sexes by downregulating the transcription in the homogametic sex, either across an entire chromosome or on a gene-by-gene basis^1,4,5^. Mechanisms of dosage compensation have been particularly well studied in male heterogametic systems (i.e. mammals), where the transcription of an entire X chromosome in females is silenced, known as X chromosome inactivation. Alternatively, the expression of X chromosome-linked genes is subjected to sex-specific regulation^6^. Dosage compensation mechanisms are largely unexplored in species with female heterogametic systems (e.g., birds), although silencing of an entire sex chromosome in males as in mammals is viewed as unlikely^7– 10^, because Z-linked genes are expressed 1.5 - 1.7 times more in males^10,14–16^. In complete dosage compensation the mean male-to-female expression ratio would be 1, while in complete absence of compensation it would be 2.

A potential mechanism regulating sex-specific dosage compensation is DNA methylation (DNAm), an epigenetic modification mostly found in CpG dinucleotides (cytosine 5′ to a guanine, separated by a phosphate bond) which suppresses gene expression^11^. Elevated levels of DNAm downregulate transcription, thereby contributing to gene inactivation. For example, an increase in the degree of DNAm of the majority of CpG islands (DNA sequences with high CG content and frequency of CpG dinucleotides) of inactivated X chromosome gene promoters and intergenic CpG islands was observed in X chromosome inactivation^12^. In contrast, the DNAm level of the avian Z chromosome was found to be similar between the sexes in chicken (*Gallus gallus*) and white-throated sparrow (*Zonotrichia albicollis*)^13^, suggesting absence of dosage compensation through DNAm in birds^10,14–17^. However, similar overall levels of DNAm do not exclude partial, gene-specific silencing, which in some avian species occurs in highly methylated regions, also known as dosage compensation valleys. Specifically, in male fowl (e.g., peacock *Pavo cristatus* and chicken *Gallus gallus domesticus*^15,20,21^), dosage compensation valleys on the Z chromosomes are known as male hypermethylated regions (MHMs)^13,18,19^. MHMs increase chromatin accessibility in females relative to males^13,17^, inducing dosage compensation of genes in the proximity of MHMs^20^. However, MHMs have been lost in passerines^14,16,10,22^ and ratites^16^, and the role of sex chromosome DNAm in dosage compensation mechanism(s) remains largely unknown for the vast majority of bird species.

If DNAm is the main mechanism driving avian dosage compensation, this should be evident as an elevated mean and variance in DNAm levels along the male Z chromosome compared to the female Z chromosome, in particular in gene promoter regions where DNAm is known to block gene expression. On the other hand, genes with detrimental effects on females are overrepresented on the Z chromosome (e.g., genes enhancing male reproductive function at the cost of females), and regulatory mechanisms have evolved to reduce their expression in females^23^. Thus, DNAm may be important in neutralizing intra-genomic sexual conflict. Moreover, genes with a female-biased transcription have mostly been translocated *away from*, rather than *to*, the Z chromosome^24^. Both processes would result in increased DNAm level on the female Z chromosome. Thus, there are opposing selection pressures on Z chromosome methylation, with an unknown net outcome on overall DNAm levels.

Due to its sex-specific inheritance, the W chromosome is subjected to natural selection that enhances female fitness^25^, leading us to expect low DNAm of the W chromosome, given that DNAm would suppress expression of W based fitness enhancing genes (e.g., related to egg-laying). However, another unique feature of the avian W chromosome, is the high frequency of endogenous retroviruses^26^, many at full length and hence potentially active, likely resulting in an elevated mutational load in females. Increased levels of DNAm on the W chromosome could potentially supress such retroviral activity^27^. Thus, there are opposing selection pressures on the W chromosome methylation, and little is known of the net outcome, except that the W chromosome is highly methylated in chicken^28,29^.

In conclusion, there are opposing evolutionary forces driving sex-specific DNAm levels on both sex chromosomes, the net outcome of which can only be revealed empirically. We examined DNAm levels in two passerine birds: captive zebra finches *Taeniopygia guttata* and wild jackdaws *Corvus monedula*. Specifically, we compared DNAm on the chromosome level of autosomes and sex chromosomes and the distribution of hypo- and hyper-methylated sites over different functional genomic regions between the sexes, with the aim to better understand sex chromosome evolution facilitated by dosage compensation and the role of the differences in shaping sexual phenotypes.

## Materials and methods

### Samples

Exact age at sampling of all individuals was known because subjects were followed since birth in both species^30,31^. Zebra finch samples were collected in the context of a long-term experiment in outdoor aviaries (320 × 150 × 210 cm) each containing single-sex flocks with 18–24 adults^32^. Selected blood samples were taken from ten individuals (six males and four females), each sampled twice (20 genomes) with an average interval of 1,470 days, between June 2008 and December 2014. Average age at sampling was 464 days (SD: 237.0) and 1,934 days (SD: 767.2) at collection of the first and second blood sample, respectively (see Table S1 for exact ages). Jackdaws were sampled in the context of a long-term study^33^ of a free-ranging population breeding in nest-boxes south of Groningen, the Netherlands (53.1708°N, 6.6064°E). For the present study, we selected 22 blood samples (genomes) collected from 11 known age adults (five males and six females); all individuals were sampled twice with an average interval at 2,429 days in the years 2007 to 2021. The average age at sampling was 877 days (SD: 241.2) and 3,306 days (SD: 947.9) during collection of the first and second samples, respectively (Table S1).

### Whole genome bisulphite sequencing

We extracted DNA according to the manufacturer’s protocol using innuPREP DNA Mini Kit (Analytik Jena GmBH) from 3 uL (nucleated) red blood cells stored in glycerol storage buffer (40% glycerol, 50mM TRIS, 5mM MgCl_2_, 0.1mM EDTA) at -80°C. Next-generation sequencing was outsourced to The Hospital for Sick Children (Toronto, Canada), where paired-end Illumina next-generation sequencing (150bp) was carried out on either an Illumina HiSeqX™ (12 zebra finch samples) or an Illumina NovaSeq™ S4 flowcell sequencer (eight zebra finch samples and 22 jackdaw samples). Libraries were prepared using the Swift Biosciences Inc. Accel NGS Methyl Seq kit (part no. 30024 and 30096) and DNA was bisulphite converted using the EZ-96 DNA Methylation-Gold kit (Zymo Research Inc., part no. D5005) as per the manufacturer’s protocol and subsequently subjected to whole-genome amplification.

### Bioinformatic processing of the whole-genome bisulphite sequencing data

Sequences were trimmed using Trim Galore!^34^ in paired-end mode, while filtering for low-quality bases (Phred score < 20). Visual controls of the data for sequence qualities, duplication levels, adapter content etc., were carried out before and after trimming using FastQC^35^ and MultiQC^36^. Because the Accel-NGS Swift kit was used for library preparation, the first ∼10 bp showed extreme biases in sequence composition and M-bias, so after checking the M-bias plots, the first 10 bps were trimmed from each sequence.

Alignments were performed using Bismark v. 0.14.4^37^ using the Bowtie 2^38^ alignment algorithm for both *in silico* bisulphite conversion of the reference genomes and alignments (see codes). For zebra finch, trimmed reads were aligned against the *in silico* bisulphite converted zebra finch reference genome (bTaeGut1.4.pri^39^; https://www.ncbi.nlm.nih.gov/data-hub/genome/GCF_003957565.2/). While some bird genome assemblies contain the sex chromosomes, many reference genomes (including the current jackdaw reference genome assembly) originate from males and thus lack a W chromosome sequence. Therefore, the jackdaw sequencing data were aligned to a bisulphite converted Hawaiian crow genome (*Corvus hawaiiensis*, bCorHaw1.pri.cur^39^; https://www.ncbi.nlm.nih.gov/data-hub/genome/GCF_020740725.1/), which, is a chromosome-level Vertebrate Genomes Project^39^ genome assembly, which is annotated allowing to assess the functional consequences of DNAm levels. The Hawaiian crow is the most closely related species for which a high-quality genome assembly was available. The divergence time between the crow and jackdaw clades has been estimated at 13 million years^40^. To verify the robustness of our findings, we repeated the analyses aligning the jackdaw sequences to the bisulphite converted jackdaw reference genome (Table S2) along with the mitochondrial alignment.

DNAm calling was performed using the Bismark methylation extractor of the Bismark Bisulfite Mapper^37^. We selected Bismark, which uses the Bowtie 2 alignment algorithm as an aligner because of its integrated features and relative resistance to error across ranges of DNAm levels compared to other packages^41^. We obtained mean DNAm percentage and its standard deviation per chromosome per sample, averaged over all CpG sites. Mapping efficiencies were 64.5 % (SD: 3.22) and 64.7 % (SD: 0.98) for the zebra finch and jackdaw respectively.

### Methylation annotation

The files containing DNAm level information for each sample were merged and used to identify hypo- and hypermethylated sites across all samples. We calculated the average DNAm percentage for each CpG site separately for the sexes, and weighted by sample dependent coverage of that site. Based on these averages, we scored sites which were present in all samples as being either hypomethylated (<= 10% on average, enhanced potential for transcription), hypermethylated (>= 90% on average, reduced transcription). We then combined the location of these sites with a functional annotation of the genomes. The annotation files for the zebra finch (GCF_003957565.2) and the jack-down (GCF_020740725.1) reference genomes were retrieved in gene transfer format (GTF). These GTF annotation files were then converted into BED12 format using the University of California, Santa Cruz (UCSC) utilities gtfToGenePred and genePredToBed (available at https://hgdownload.soe.ucsc.edu/downloads.html#utilities_downloads). CpG sites were annotated using annotateWithGeneParts from the R 4.1.2 package Genomation 1.4.1^42^. This tool hierarchically classifies sites into pre-defined functional regions, i.e., promoter, exon, intron, or intergenic. The pre-defined functional regions were based on the annotation information present in the BED12 files accessed with the Genomation tool readTranscriptFeatures. A customized script was employed to integrate the annotation results of CpG sites with their respective gene symbol information.

### Statistical analyses

As dependent variables, we used chromosome-specific mean and standard deviation of the DNAm level in each sample, and assessed these data using linear mixed models (LMMs) in R version 4.2.2^43^. The fixed effects considered were chromosome type (autosome, W and Z), sex (male vs. female), age (young = first sample vs. old = second sample per individual) and chromosome length (in base pairs, bp, natural log-transformed). Chromosome number, sample ID, and individual ID were included as random effects. To facilitate interpretation of the statistical output, we created dummy variables^44^ for chromosome type as follows: variable ‘W’ was coded as 1 for W and 0 for both other types, and analogously for Z. Sex and age were also coded as dummy variables. We mean-centred all fixed effects to facilitate the interpretation of the estimates. We tested two-way interactions, and sequentially removed the least significant terms until all remaining variables were either statistically significant or part of significant interactions. We calculated *R*^2^ as defined in^45^ for each model with the MuMIn R package^46^. We tested all models for multicollinearity (VIF scores never exceed 1.1). The proportion of hypo- or hypermethylated CpG sites in each annotation category (promoters, exons, introns and intergenic regions) was compared with the expected proportion for each chromosome type (autosomes, W and Z) using an exact binomial test. The proportion of expected CpG sites per annotation category was estimated as the proportion of all CpG sites captured by our analysis located in each annotation category. To compare differences in the proportion of hypo- or hypermethylated CpG sites in each annotation category between males and females for autosomes and Z we used two-sample tests for equality of proportions in the *stats*^43^ package.

## Results

Unless otherwise mentioned, the results were qualitatively similar in both study species. DNAm level on the autosomes was similar between the sexes, but the level of DNAm on the Z chromosome was higher in females compared to males (Fig. 1; adjusting for chromosome length, Table 1). This “sex” effect was of a similar magnitude in both species, approaching significance in zebra finches (P = 0.054), and significant in jackdaws (*P* = 0.014; Table 1A). Furthermore, the Z chromosome level of DNAm was significantly higher than the levels observed on the autosomes in zebra finches, but not in jackdaws (Fig. 1; adjusting for chromosome length, Table 1). The variation in DNAm level, estimated from the standard deviation (SD), did not differ significantly between the Z chromosome and autosomes when both sexes were combined. However, the SD of the DNAm level along the Z chromosome differed strongly between the sexes, being higher in females (Table 1B, Fig. 2A-B).

**Table 1.**
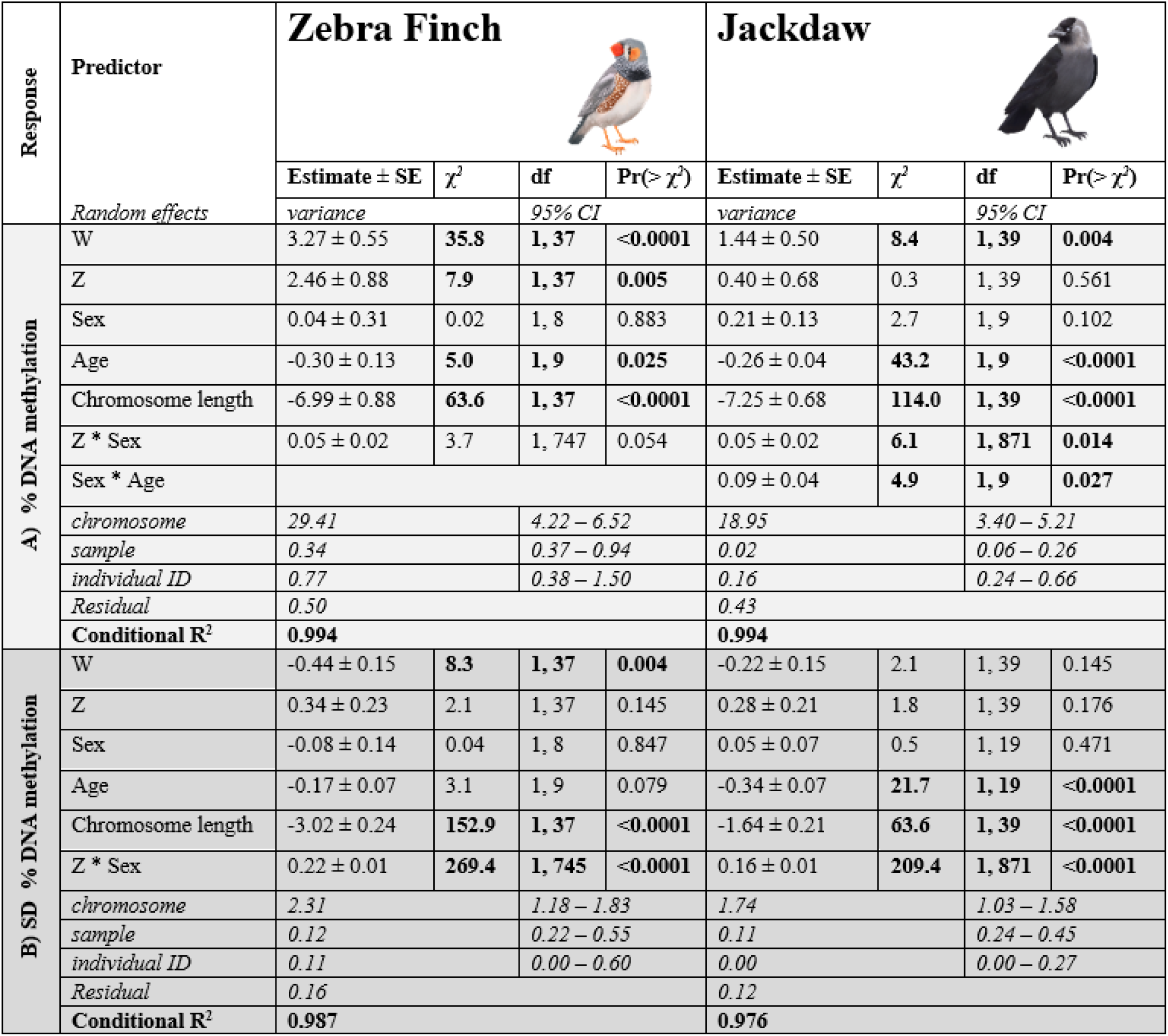
Linear mixed models examining variation of mean percentage DNA methylation (A) and standard deviation of percentage in DNA methylation (B) in zebra finches and jackdaws. Estimates are shown after mean centring of all predictors. Chromosome length was ln-transformed. Autosomes, males and young individuals were the reference categories for predictors. We present the final models with nonsignificant (P < 0.05) interactions removed. Sample sizes for each analysis are shown in Table S1. Significant differences (P < 0.05) are indicated in bold.

**Figure 1.**
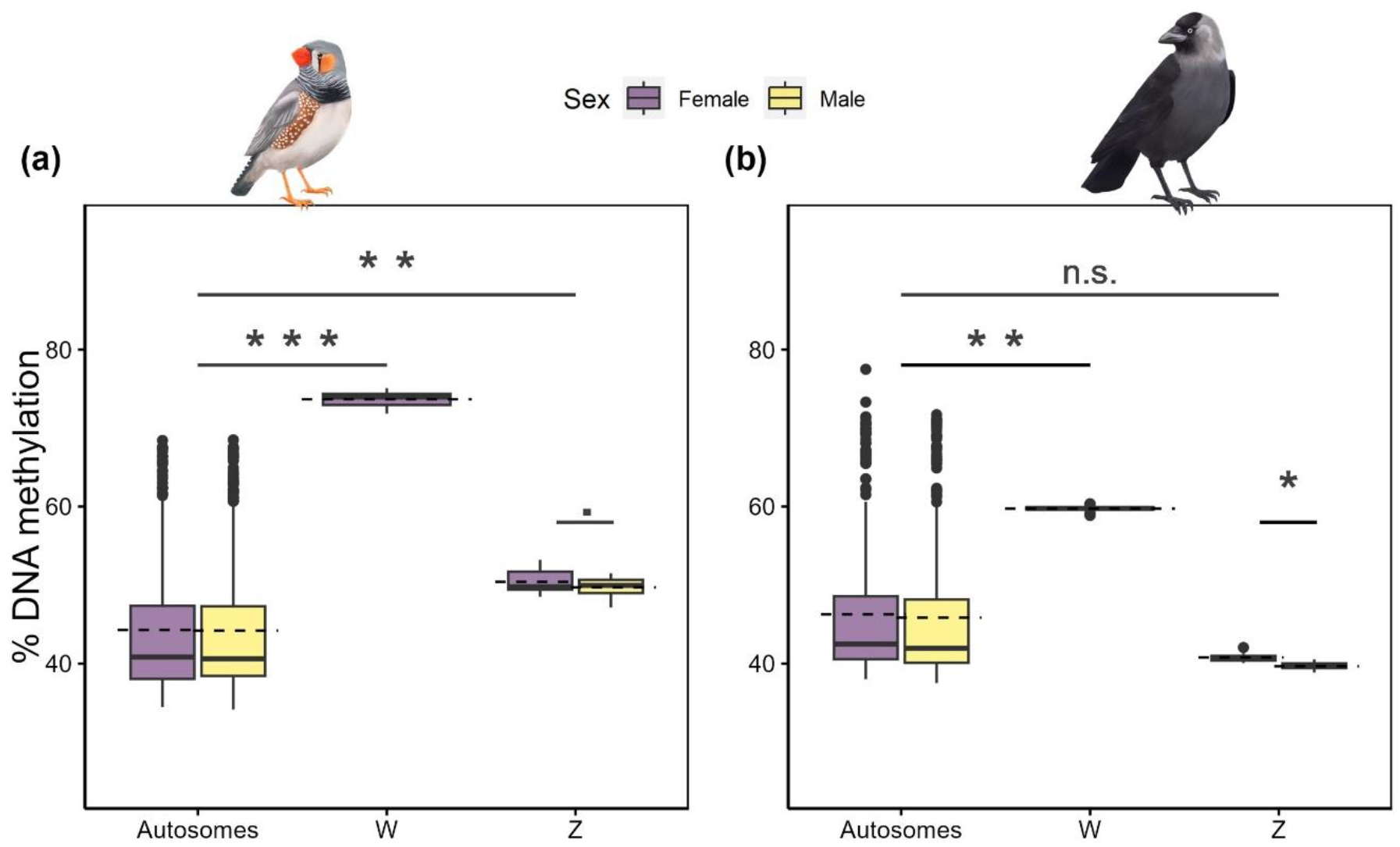
Chromosome type specific DNAm in male and female (a) zebra finches and (b) jackdaws. Boxplots on raw data represent median values, interquartile ranges and outliers (dots) and dashed lines represent the means. Estimates shown were based on the average DNAm per chromosome in each sample. Significance from Table 1 (controlling for chromosome length) indicated by: *** p < 0.001, ** p < 0.01, * p < 0.05, n.s. p > 0.05.

**Figure 2.**
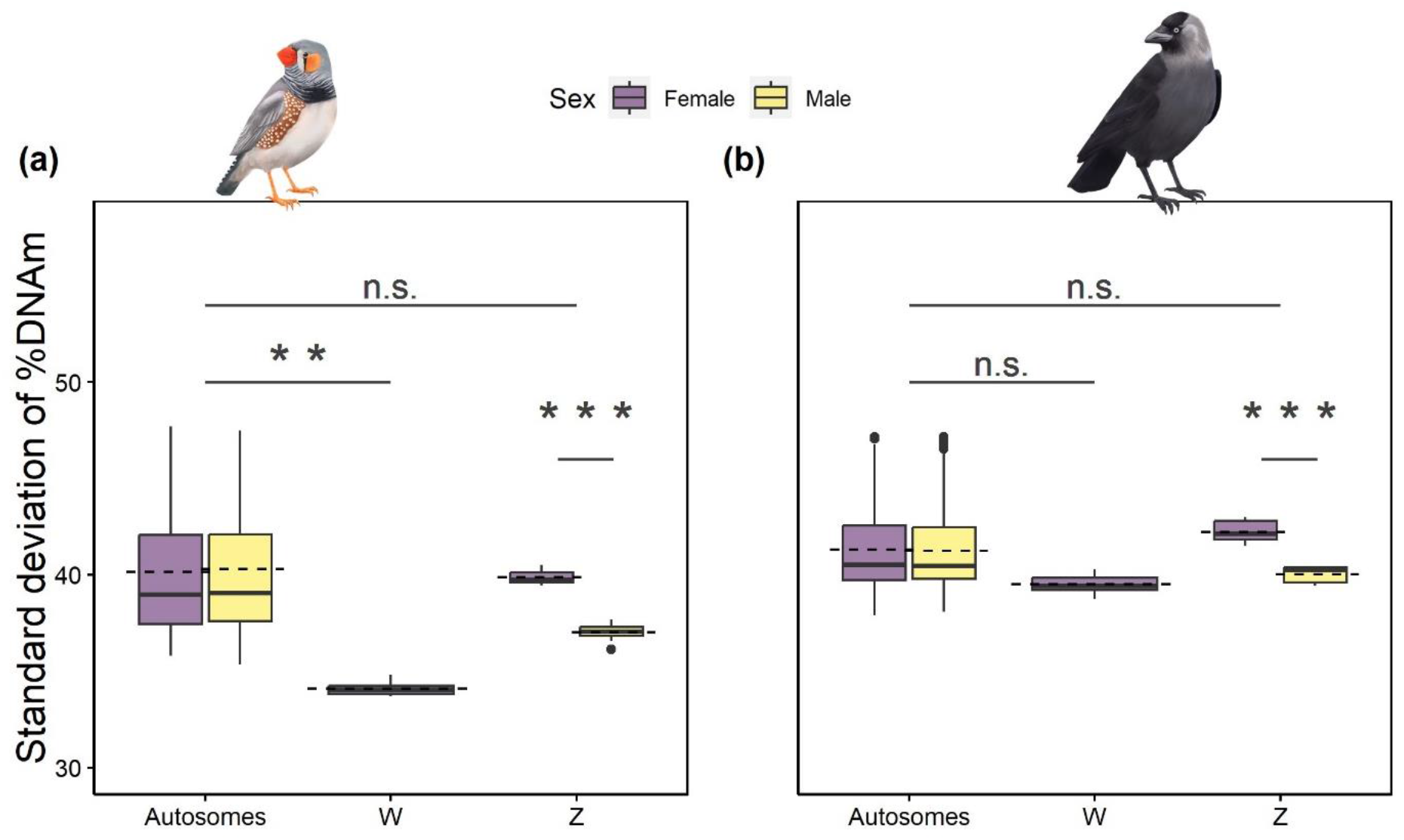
The standard deviation of the percentage of DNAm per chromosome in (a) zebra finches and (b) jackdaws depending on the chromosome type and sex. Boxplots on raw data represent median values, interquartile ranges, and outliers (dots) and dashed lines represent the mean. Significance from Table 1 (controlling for chromosome length) indicated by: *** p < 0.001, ** p < 0.01, n.s. p > 0.05.

Across all chromosomes, the proportion of hypomethylated sites (DNAm <= 10%) in gene promoter regions was higher than expected by chance, and, conversely, the proportion of hypermethylated sites (DNAm >= 90%) in gene promotor regions was lower than expected by chance (Fig. 3, Table S5). The proportion of DNA hypo- and hypermethylation of promoter regions located on autosomal chromosomes in the zebra finch was higher in females, compared to males (Fig. 3, Table S4). Hypermethylated sites on Z chromosomes were predominantly located in intergenic regions and introns (Fig. 3). DNA hypermethylation on the Z chromosome was overrepresented in exons and introns and underrepresented in promoter and intergenic regions, regardless of sex (Table S5). We detected a higher degree of hypomethylation in promoter regions on the Z chromosome in males, an effect that was particularly strong in jackdaws. The level of hypermethylation in promoter regions, while overall low, was also slightly higher on the Z chromosome in jackdaw males (Fig. 3, Table S4).

**Figure 3.**
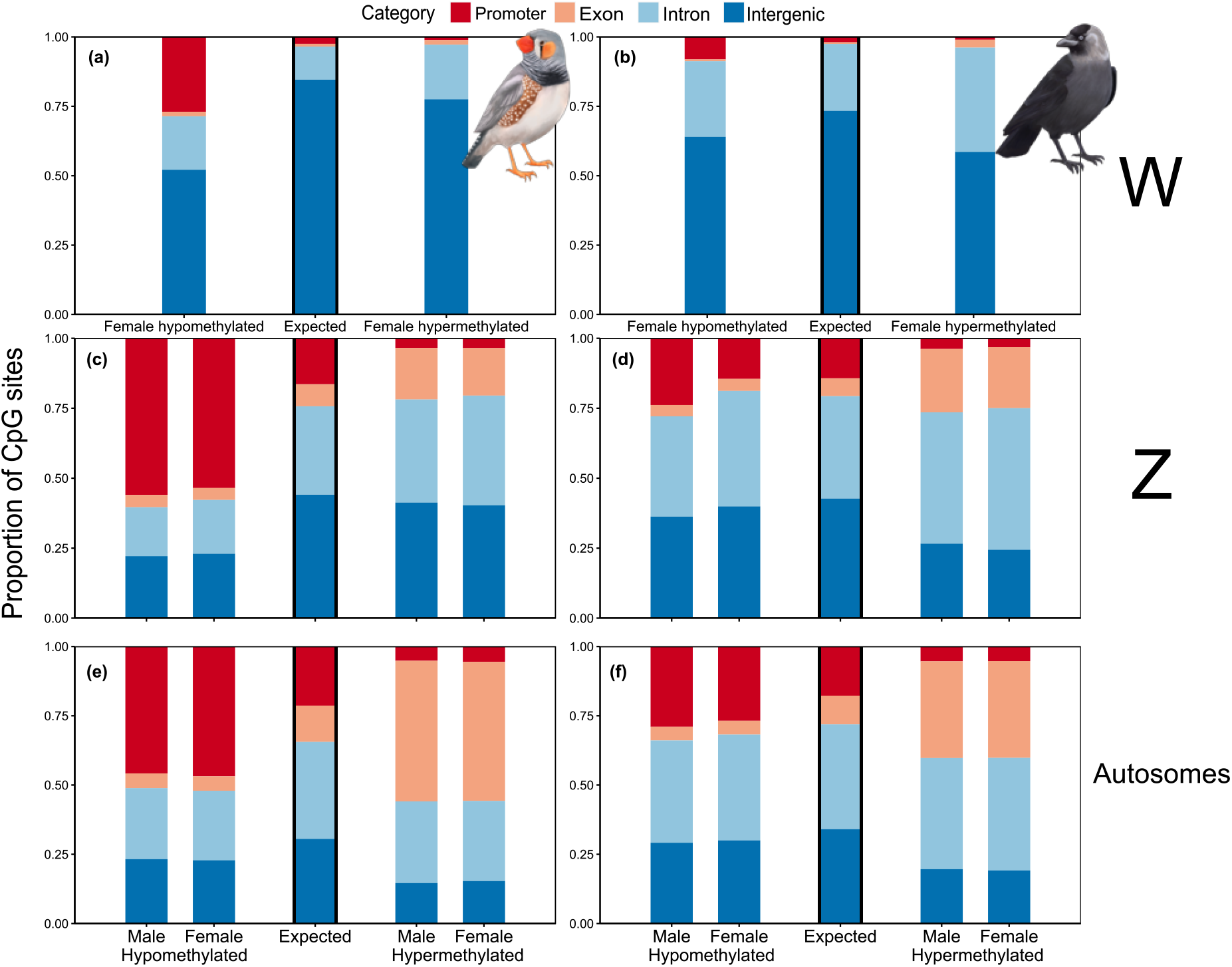
Proportion of hypomethylated (<=10%) and hypermethylated (>=90%) CpG sites across all samples on the W (a - b), Z (c - d), and all autosomes (e - f). Panels (a), (c) and (e) show data from zebra finches, and (b), (d) and (f) from the jackdaws. The order of precedence for the functional annotation categories was promoter > exon > intron. The expected proportion of CpG sites per annotation category was estimated as the proportion of all CpG sites captured by our analysis located in each annotation category.

The level of DNAm was substantially higher on the W chromosome compared to all other chromosomes (Fig. 1; adjusting for chromosome length, Table 1). In line with this finding, the proportion of hypermethylated sites was higher on the W chromosome compared to all other chromosomes (Fig. S7). Similarly to the patterns detected on the Z chromosome, hypermethylated CpG sites on the W chromosome were predominantly located in intergenic regions, followed by introns, exons and promoters (Fig. 3). Hypermethylated CpG sites on the W chromosome were overrepresented in introns and exons and underrepresented in promoter and intergenic regions (Table S5).

## Discussion

We investigated chromosome-level DNAm of autosomes and sex chromosomes in two female heterogametic species to test predictions based on the hypothesis that DNAm plays a key role in driving dosage compensation. The predictions that DNAm level should be higher and more variable on male Z chromosomes compared to female Z chromosomes, thus contributing to gene silencing, were not confirmed. In contrast, we found the Z chromosome to be more methylated in females, and the level of DNAm on the female Z to be more variable (Fig. 1 and 2, Table 1). Furthermore, functional annotation of CpG sites revealed that hypomethylated CpG sites located in gene promoter regions on the Z chromosome were more frequent in males (Fig. 3). In contrast, the proportions of DNA hypo- and hypermethylation in promotor regions on autosomal chromosomes were higher in females in the zebra finch (Fig. 3, Table S4). These findings explain the observed higher level of gene expression of genes on the avian Z chromosome in males compared to females^10,14–17^, but are inconsistent with dosage compensation through DNAm. We therefore conclude that DNAm is not the main mechanism driving dosage compensation in passerines.

Possibly, the dosage compensation mechanism in passerines is elusive because it is not complete^10,14–17^ and might play a lesser role beyond the embryonic stage. In contrast to mammals, where sex hormones are crucial for the expression of sex-specific phenotypes, avian sexual phenotypes appear little dependent on the dosage of genes regulating sex hormones. Instead, birds display a cell-autonomous sex identity, meaning that sex chromosome content of each cell determines its sexual phenotype^47^. The limited extent to which dosage compensation is achieved, given that Z chromosome-based gene expression in males is not two times higher than Z chromosome-based gene expression in females, might instead rely on microRNA regulation^48^ or other epigenetic elements such as chromatin accessibility^49^. However, this does not explain why female Z chromosomes are more methylated than male Z chromosomes. In female heterogametic systems, Z chromosome evolution favours the presence of male-beneficial alleles^24^, resulting in overrepresentation of female-detrimental genes^23^. An effort to silence those genes likely explains the higher level of DNAm on the female Z chromosome compared to males.

Expectations with respect to DNAm level of the W chromosome were mixed, because on the one hand the presence of female fitness enhancing genes should lead to a low DNAm level, while on the other hand the preponderance of transposable elements on the W chromosome^26,27^ should lead to a high level of DNAm. Evidently, the net outcome of these opposing selective pressures results in high DNAm levels on the W chromosome (Fig. 1), as previously also described in chickens^28,29,50^. In line with this inference, we detected a high proportion of hypermethylated sites in intergenic regions on the W chromosome, which is where most transposable elements are located^51^ (Fig. 3). Our expectation that genes on the W chromosome, that presumably benefit female fitness, should display low DNAm levels was however also confirmed: gene promoter regions on the W chromosome were frequently hypomethylated (Table S5), suggesting elevated transcription of W chromosome-linked genes^25^. Thus, site, or gene-specific, elevated or decreased levels of DNAm across both sex chromosomes are likely involved in the development of sex-specific phenotypes in female heterogametic systems rather than driving dosage compensation between the avian sex chromosomes. Further corroboration of DNAm as mediator of the evolution of sexually antagonistic genes^23,24^ is provided by the fact that sex-specific differences in level and variance of DNAm were limited to the Z chromosome, i.e. were absent from autosomes (Fig 1, Fig. 2).

In conclusion, we presented a chromosomal perspective of the DNAm landscape in birds that offers new insights on the role of sex chromosomes in the evolution of female heterogametic sex-determination systems. Our results prompt a plethora of key questions concerning the effects of DNAm-mediated differential gene expression and its role in shaping sex-dependent traits which impact fitness. We think of the epigenome as a continuously optimized trait, adaptively changing throughout an individual’s lifetime, to maximize fitness and survival. We therefore predict that the patterns we observed will relate to variation in life stage, underlying genomic variants, and state of the environment, and see analysis of such variation as a promising next step to further our understanding of adaptation and genome evolution.

## Acknowledgements

We thank the Center for Information Technology of the University of Groningen for their support and for providing access to the Peregrine and Hábrók high performance computing clusters. We express our gratitude to Ellis Mulder for her help in the laboratory and to Berber Maarsingh for the beautiful drawings of our study species used in the graphs of this manuscript. We also thank the animal caretakers of the University of Groningen as well as numerous students whose invaluable help made this project possible.

## Author contributions

JS, MT, PJP and SV conceived and designed the study. JS, MT and FF performed the bioinformatics and data analysis. JS wrote the manuscript with substantial input from all authors. All authors approved the final version of the manuscript.

## Ethics approval

All methods and experiments for the zebra finches detailed in this manuscript were performed under the approval of the Central Committee for Animal Experiments (Centrale Commissie Dierproeven) of the Netherlands, under licenses AVD1050020174344 and AVD1050020184967.

