## Supplementary Material for "Dosage compensation and sexual conflict in female heterogametic methylomes"

**Affiliations:**

Analysis performed on jackdaw samples aligned to bisulfite converted jackdaw genome

To align jackdaw samples we used reference genome of *Corvus monedula*, ASM1340703v1[1]; https://www.ncbi.nlm.nih.gov/data-hub/genome/GCA_013407035.1/, for which the mapping efficiency was 76.5% (±SD: 0.97). The analysis performed on jackdaw samples aligned to bisulphite converted jackdaw genome confirmed that average DNA methylation (DNAm) decreased with age (Table S2A). Accordingly with jackdaw samples aligned to the more detailed Hawaiian crow genome, there were no sex specific differences but a significant interaction of chromosome type and sex revealed that Z chromosome was more methylated in females than in males (Fig. S2). The main differences was that average DNAm level of Z chromosome was higher compared to autosomes (Fig. S3, but note that in jackdaw aligned to jackdaw genome there are no microchromosomes and W assembled that are all short and highly methylated, Fig. S4). For the same reason we detected no association of average DNAm level with chromosome length (Table S3A). Nevertheless, the absolute values for DNAm level were higher in jackdaw aligned to jackdaw genome than in jackdaw aligned to Hawaiian crow genome (Figs. S3-S4). This may stem from the fact that sequences that aligned to W and microchromosomes in the Hawaiian crow might have been included in the alignment to Z and other autosomes in jackdaw genome. Standard deviation (SD) of percentage in DNAm level was uniform between Z chromosome and autosomes (Table S2B, Fig. S5). SD was negatively associated with chromosome length and we detected lower values in old individuals compared to young (Table S2B), consistently with the relation observed for average DNAm level values. There was no difference in SD DNAm level between sexes, but we detected a significant interaction showing that Z chromosome was more variably methylated in females than males (Table S2B, Fig. S5), a much stronger effect while compared to jackdaws aligned to HC genome. Additionally, a significant interaction of chromosome type and age showed that the decrease in SD DNAm level with age was higher in autosomes than in Z chromosome (Table S2B). SD in jackdaw aligned to jackdaw genome was in general lower than when aligned to Hawaiian crow (Fig. S6). Inspiringly, our findings on the absence of DNA-mediated dosage compensation are robust, regardless of which reference genome was used in the case of the jackdaws. We show that sex-specific DNAm level patterns were largely consistent between the two reference genomes (complete Hawaiian crow and male only jackdaw) used for aligning jackdaw samples (Supplementary Material, Table S2, Fig. S2-S6).

Mitochondrial DNA methylation

Mitochondrial DNA (mtDNA) methylation was invariably low in both sexes and species (overall averages: 0.7%, SD: 0.37 in zebra finches; 0.5%, SD: 0.40 in jackdaws). mtDNA average methylation

level and SD were higher in females compared to males (Table S3, separate analysis in both species combined).
mtDNA methylation level, although overall low, was higher and more variable in females than in males (Table S3). These very low levels are congruent with studies in other taxa[2], however there is a debate around the possibilities of detection of mtDNA methylation with studies in mice and humans pointing to absence of mtDNA methylation[3,4] or DNAm present in other than CpG context[5]. It is not necessarily the case in birds as they have lower mtDNA variability than mammals possibly because both W and mtDNA are maternally transmitted and as such do not recombine[6]. It is hard to speculate on the function of the detected sex-specific mtDNA methylation levels since not much is known about functional roles of mtDNA methylation in general[7].

Table S1. Zebra finch and jackdaw longitudinal sample information

| Species | BirdID | Age young (days) | Age old (days) | Sex |
|---|---|---|---|---|
| **Zebra finches** | 2269 | 295 | 2636 | male |
| | 2288 | 289 | 2644 | female |
| | 2257 | 298 | 884 | male |
| | 2277 | 292 | 631 | male |
| 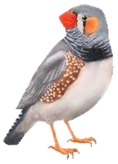 | 2247 | 468 | 2651 | female |
| | 1923 | 853 | 2124 | male |
| | 2172 | 732 | 2002 | female |
| | 1942 | 847 | 2484 | male |
| | 2354 | 351 | 2367 | male |
| | 2423 | 213 | 912 | female |
| **Jackdaws** | 480 | 749 | 5496 | female |
| | 775 | 743 | 3666 | male |
| | 888 | 748 | 2584 | female |
| 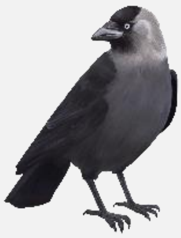 | 923 | 753 | 2948 | female |
| | 973 | 742 | 3670 | male |
| | 977 | 1119 | 3666 | male |
| | 1146 | 1458 | 4022 | female |
| | 1350 | 1099 | 2939 | male |
| | 1502 | 745 | 2935 | female |
| | 1605 | 747 | 2228 | female |
| | 1762 | 745 | 2210 | male |

Table S2. Linear mixed models examining variation of mean percentage DNA methylation (A) and standard deviation of percentage in DNA methylation (B) in jackdaws aligned to jackdaw genome. Estimates are shown after mean centering of all predictors. Chromosome length was log transformed. Autosomes, males and young individuals were the reference categories for predictors. We present the final models with nonsignificant interactions removed. Sample sizes for each analysis are shown in Table S1. Significant differences (P < 0.05) are indicated in bold.

| **Response** | **Predictor** | **Jackdaw** 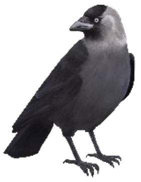 | | | |
|---|---|---|---|---|---|
| | | **Estimate ± SE** | **$\chi^2$** | **df** | **Pr(> $\chi^2$)** |
| | *Random effects* | *variance* | | *95% confidence interval* | |
| **A) % DNA methylation** | Z | 0.86 ± 0.32 | **7.3** | **1, 26** | **0.007** |
| | Sex | 0.26 ± 0.19 | 1.9 | 1, 9 | 0.164 |
| | Age | -0.31 ± 0.06 | **27.5** | **1, 10** | **<0.0001** |
| | Chromosome length | 0.06 ± 0.32 | 0.03 | 1, 26 | 0.864 |
| | Z * Sex | 0.04 ± 0.01 | **30.7** | **1, 587** | **<0.0001** |
| | *chromosome* | *2.74* | | *1.24 – 2.09* | |
| | *sample* | *0.08* | | *0.18 – 0.42* | |
| | *individual ID* | *0.35* | | *0.35 – 0.94* | |
| | *Residual* | *0.03* | | | |
| | **Conditional $R^2$** | **0.992** | | | |
| **B) SD DNA methylation** | Z | 0.27 ± 0.14 | 3.6 | 1, 26 | 0.057 |
| | Sex | 0.05 ± 0.09 | 0.3 | 1, 19 | 0.574 |
| | Age | -0.43 ± 0.09 | **22.4** | **1, 19** | **<0.0001** |
| | Chromosome length | -1.22 ± 0.14 | **75.2** | **1, 26** | **<0.0001** |
| | Z * Sex | 0.22 ± 0.01 | **1300.6** | **1, 586** | **<0.0001** |
| | Z * Age | 0.01 ± 0.01 | **5.4** | **1, 586** | **0.021** |
| | *chromosome* | *0.54* | | *0.55 – 0.92* | |
| | *sample* | *0.18* | | *0.30 – 0.56* | |
| | *individual ID* | *<0.0001* | | *0.00 – 0.35* | |
| | *Residual* | *0.02* | | | |
| | **Conditional $R^2$** | **0.990** | | | |

Table S3. Linear mixed models examining variation of mean percentage DNA methylation (A) and standard deviation of percentage in DNA methylation (B) in mitochondrial DNA in zebra finches and jackdaws combined. Estimates are shown after mean centering of sex. Males and jackdaws were the reference categories for predictors. We present the final models with nonsignificant interactions removed. Sample sizes for each analysis are shown in Table S1. Significant differences ($P < 0.05$) are indicated in bold.

| **Response** | **Predictor** / *Random effect* | **Estimate ± SE** / *variance* | **$\chi^2$** | **df** / *95% confidence interval* | **Pr(> $\chi^2$)** |
|---|---|---|---|---|---|
| **A) % DNA methylation** | Sex | 0.18 ± 0.07 | **7.3** | **1, 18** | **0.007** |
| | Species | 0.25 ± 0.13 | 3.5 | 1, 18 | 0.062 |
| | *individual ID* | *0.06* | | *0.00 – 0.34* | |
| | *Residual* | *0.07* | | | |
| | **Conditional $R^2$** | **0.585** | | | |
| **B) SD DNA methylation** | Sex | 1.36 ± 0.44 | **9.3** | **1, 18** | **0.002** |
| | Species | 1.48 ± 0.88 | 2.8 | 1, 18 | 0.093 |
| | *individual ID* | *1.66* | | *0.00 – 2.13* | |
| | *Residual* | *4.64* | | | |
| | **Conditional $R^2$** | **0.449** | | | |

Table S4. Proportions of distributions of CpG sites in different annotation categories in the two sexes in zebra finches (a) and jackdaws (b), with average DNAm level per site across all samples being hypomethylated (<=10%) and hypermethylated (>=90%). Shown for autosomes and Z; results of two-sample test for equality of proportions between males vs. females in each annotation category. Significant differences (P < 0.05) are indicated in bold.

**males vs. females proportions of hypo- and hypermethylated sites in zebra finch autosomes and Z**

**(a)**

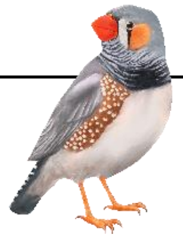

| category | | Promoters | | Exons | | Introns | | Intergenic | |
|---|---|---|---|---|---|---|---|---|---|
| **Autosomes** | **DNAm** | males vs. females | | males vs. females | | males vs. females | | males vs. females | |
| | ***Hypo*** | Proportion males | Proportion females | Proportion males | Proportion females | Proportion males | Proportion females | Proportion males | Proportion females |
| | | 0.458 | 0.469 | 0.053 | 0.052 | 0.256 | 0.251 | 0.232 | 0.228 |
| | | **<0.0001** | | **<0.0001** | | **<0.0001** | | **<0.0001** | |
| | ***Hyper*** | Proportion males | Proportion females | Proportion males | Proportion females | Proportion males | Proportion females | Proportion males | Proportion females |
| | | 0.051 | 0.055 | 0.508 | 0.502 | 0.295 | 0.289 | 0.146 | 0.153 |
| | | **<0.0001** | | **<0.0001** | | **<0.0001** | | **<0.0001** | |
| **Z** | **DNAm** | males vs. females | | males vs. females | | males vs. females | | males vs. females | |
| | ***Hypo*** | Proportion males | Proportion females | Proportion males | Proportion females | Proportion males | Proportion females | Proportion males | Proportion females |
| | | 0.560 | 0.535 | 0.044 | 0.042 | 0.175 | 0.192 | 0.221 | 0.230 |
| | | **<0.0001** | | 0.995 | | **<0.0001** | | 0.995 | |
| | ***Hyper*** | Proportion males | Proportion females | Proportion males | Proportion females | Proportion males | Proportion females | Proportion males | Proportion females |
| | | 0.034 | 0.035 | 0.184 | 0.170 | 0.369 | 0.392 | 0.413 | 0.403 |
| | | 0.584 | | **<0.0001** | | **<0.0001** | | **0.002** | |

**males vs. females proportions of hypo- and hypermethylated sites in jackdaw autosomes and Z**

# (b)

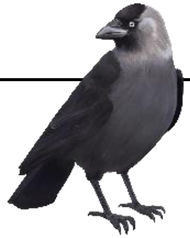

| category | | Promoters | | Exons | | Introns | | Intergenic | |
|---|---|---|---|---|---|---|---|---|---|
| **Autosomes** | **DNAm** | males vs. females | | males vs. females | | males vs. females | | males vs. females | |
| | ***Hypo*** | Proportion males | Proportion females | Proportion males | Proportion females | Proportion females | Proportion females | Proportion males | Proportion females |
| | | 0.290 | 0.268 | 0.049 | 0.050 | 0.369 | 0.382 | 0.291 | 0.300 |
| | | **<0.0001** | | **0.020** | | **<0.0001** | | **<0.0001** | |
| | ***Hyper*** | Proportion males | Proportion females | Proportion males | Proportion females | Proportion males | Proportion females | Proportion males | Proportion females |
| | | 0.053 | 0.052 | 0.349 | 0.349 | 0.401 | 0.407 | 0.196 | 0.192 |
| | | **0.025** | | 0.891 | | **<0.0001** | | **<0.0001** | |
| **Z** | **DNAm** | males vs. females | | males vs. females | | males vs. females | | males vs. females | |
| | ***Hypo*** | Proportion males | Proportion females | Proportion males | Proportion females | Proportion males | Proportion females | Proportion males | Proportion females |
| | | 0.238 | 0.145 | 0.041 | 0.043 | 0.358 | 0.413 | 0.363 | 0.400 |
| | | **<0.0001** | | **<0.0001** | | **<0.0001** | | **<0.0001** | |
| | ***Hyper*** | Proportion males | Proportion females | Proportion males | Proportion females | Proportion males | Proportion females | Proportion males | Proportion females |
| | | 0.038 | 0.031 | 0.226 | 0.218 | 0.470 | 0.506 | 0.266 | 0.245 |
| | | **<0.0001** | | **0.001** | | **<0.0001** | | **<0.0001** | |

Table S5. Proportions of observed and expected distributions of CpG sites in different annotation categories in zebra finches (a) and jackdaws (b), with average DNAm level per site across all samples being hypomethylated (<=10%) and hypermethylated (>=90%). Shown for autosomes, Z and W; results of two-sided exact binomial tests between observed vs. expected proportions in each annotation category. Expected proportions were calculated from the location of all CpG sites captured by our analysis. Significant differences (P < 0.05) are indicated in bold.

**observed vs. expected proportions of hypo- and hypermethylated sites per chromosome type per sex in zebra finches**

**(a)**

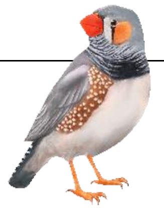

| **category** | | **Promoters** | | | | **Exons** | | | | **Introns** | | | | **Intergenic** | | | |
|---|---|---|---|---|---|---|---|---|---|---|---|---|---|---|---|---|---|
| **Autosomes** | **DNAm** | Males | | Females | | Males | | Females | | Males | | Females | | Males | | Females | |
| | | observed | expected | observed | expected | observed | expected | observed | expected | observed | expected | observed | expected | observed | expected | observed | expected |
| | ***Hypo*** | 0.458 | 0.213 | 0.469 | 0.213 | 0.053 | 0.131 | 0.052 | 0.131 | 0.256 | 0.350 | 0.251 | 0.350 | 0.232 | 0.306 | 0.228 | 0.306 |
| | | **<0.0001** | | **<0.0001** | | **<0.0001** | | **<0.0001** | | **<0.0001** | | **<0.0001** | | **<0.0001** | | **<0.0001** | |
| | | observed | expected | observed | expected | observed | expected | observed | expected | observed | expected | observed | expected | observed | expected | observed | expected |
| | ***Hyper*** | 0.051 | 0.213 | 0.055 | 0.213 | 0.508 | 0.131 | 0.502 | 0.131 | 0.295 | 0.350 | 0.289 | 0.350 | 0.146 | 0.306 | 0.153 | 0.306 |
| | | **<0.0001** | | **<0.0001** | | **<0.0001** | | **<0.0001** | | **<0.0001** | | **<0.0001** | | **<0.0001** | | **<0.0001** | |
| **Z** | **DNAm** | Males | | Females | | Males | | Females | | Males | | Females | | Males | | Females | |
| | | observed | expected | observed | expected | observed | expected | observed | expected | observed | expected | observed | expected | observed | expected | observed | expected |
| | ***Hypo*** | 0.560 | 0.164 | 0.535 | 0.164 | 0.044 | 0.079 | 0.042 | 0.079 | 0.175 | 0.315 | 0.192 | 0.315 | 0.221 | 0.441 | 0.230 | 0.441 |
| | | **<0.0001** | | **<0.0001** | | **<0.0001** | | **<0.0001** | | **<0.0001** | | **<0.0001** | | **<0.0001** | | **<0.0001** | |
| | | observed | expected | observed | expected | observed | expected | observed | expected | observed | expected | observed | expected | observed | expected | observed | expected |
| | ***Hyper*** | 0.034 | 0.164 | 0.035 | 0.164 | 0.184 | 0.079 | 0.170 | 0.079 | 0.369 | 0.315 | 0.392 | 0.315 | 0.413 | 0.441 | 0.403 | 0.441 |
| | | **<0.0001** | | **<0.0001** | | **<0.0001** | | **<0.0001** | | **<0.0001** | | **<0.0001** | | **<0.0001** | | **<0.0001** | |
| **W** | **DNAm** | Females | | | | Females | | | | Females | | | | Females | | | |
| | | observed | | expected | | observed | | expected | | observed | | expected | | observed | | expected | |
| | ***Hypo*** | 0.270 | | 0.025 | | 0.016 | | 0.010 | | 0.192 | | 0.118 | | 0.521 | | 0.847 | |
| | | **<0.0001** | | | | **<0.0001** | | | | **<0.0001** | | | | **<0.0001** | | | |
| | | observed | | expected | | observed | | expected | | observed | | expected | | observed | | expected | |
| | ***Hyper*** | 0.011 | | 0.025 | | 0.017 | | 0.010 | | 0.197 | | 0.118 | | 0.775 | | 0.847 | |
| | | **<0.0001** | | | | **<0.0001** | | | | **<0.0001** | | | | **<0.0001** | | | |

**observed vs. expected proportions of hypo- and hypermethylated sites per chromosome type per sex in jackdaws**

# (b)

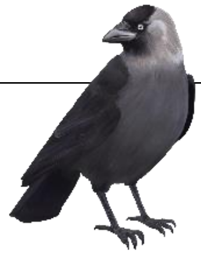

| category | | Promoters | | | | Exons | | | | Introns | | | | Intergenic | | | |
|---|---|---|---|---|---|---|---|---|---|---|---|---|---|---|---|---|---|
| **Autosomes** | **DNAm** | Males | | Females | | Males | | Females | | Males | | Females | | Males | | Females | |
| | ***Hypo*** | observed | expected | observed | expected | observed | expected | observed | expected | observed | expected | observed | expected | observed | expected | observed | expected |
| | | 0.290 | 0.177 | 0.268 | 0.177 | 0.049 | 0.104 | 0.050 | 0.104 | 0.369 | 0.377 | 0.382 | 0.377 | 0.291 | 0.341 | 0.300 | 0.341 |
| | | **<0.0001** | | **<0.0001** | | **<0.0001** | | **<0.0001** | | **<0.0001** | | **<0.0001** | | **<0.0001** | | **<0.0001** | |
| | ***Hyper*** | observed | expected | observed | expected | observed | expected | observed | expected | observed | expected | observed | expected | observed | expected | observed | expected |
| | | 0.053 | 0.177 | 0.052 | 0.177 | 0.349 | 0.104 | 0.349 | 0.104 | 0.401 | 0.377 | 0.407 | 0.377 | 0.196 | 0.341 | 0.192 | 0.341 |
| | | **<0.0001** | | **<0.0001** | | **<0.0001** | | **<0.0001** | | **<0.0001** | | **<0.0001** | | **<0.0001** | | **<0.0001** | |
| **Z** | **DNAm** | Males | | Females | | Males | | Females | | Males | | Females | | Males | | Females | |
| | ***Hypo*** | observed | expected | observed | expected | observed | expected | observed | expected | observed | expected | observed | expected | observed | expected | observed | expected |
| | | 0.238 | 0.143 | 0.145 | 0.143 | 0.041 | 0.064 | 0.043 | 0.064 | 0.358 | 0.365 | 0.413 | 0.365 | 0.363 | 0.428 | 0.400 | 0.428 |
| | | **<0.0001** | | **0.004** | | **<0.0001** | | **<0.0001** | | **<0.0001** | | **<0.0001** | | **<0.0001** | | **<0.0001** | |
| | ***Hyper*** | observed | expected | observed | expected | observed | expected | observed | expected | observed | expected | observed | expected | observed | expected | observed | expected |
| | | 0.038 | 0.143 | 0.032 | 0.143 | 0.226 | 0.064 | 0.218 | 0.064 | 0.470 | 0.365 | 0.506 | 0.365 | 0.266 | 0.428 | 0.245 | 0.428 |
| | | **<0.0001** | | **<0.0001** | | **<0.0001** | | **<0.0001** | | **<0.0001** | | **<0.0001** | | **<0.0001** | | **<0.0001** | |
| **W** | **DNAm** | Females | | | | Females | | | | Females | | | | Females | | | |
| | ***Hypo*** | observed | | expected | | observed | | expected | | observed | | expected | | observed | | expected | |
| | | 0.081 | | 0.019 | | 0.007 | | 0.007 | | 0.272 | | 0.240 | | 0.640 | | 0.734 | |
| | | **<0.0001** | | | | 0.901 | | | | **<0.0001** | | | | **<0.0001** | | | |
| | ***Hyper*** | observed | | expected | | observed | | expected | | observed | | expected | | observed | | expected | |
| | | 0.010 | | 0.019 | | 0.028 | | 0.007 | | 0.376 | | 0.240 | | 0.586 | | 0.734 | |
| | | **<0.0001** | | | | **<0.0001** | | | | **<0.0001** | | | | **<0.0001** | | | |

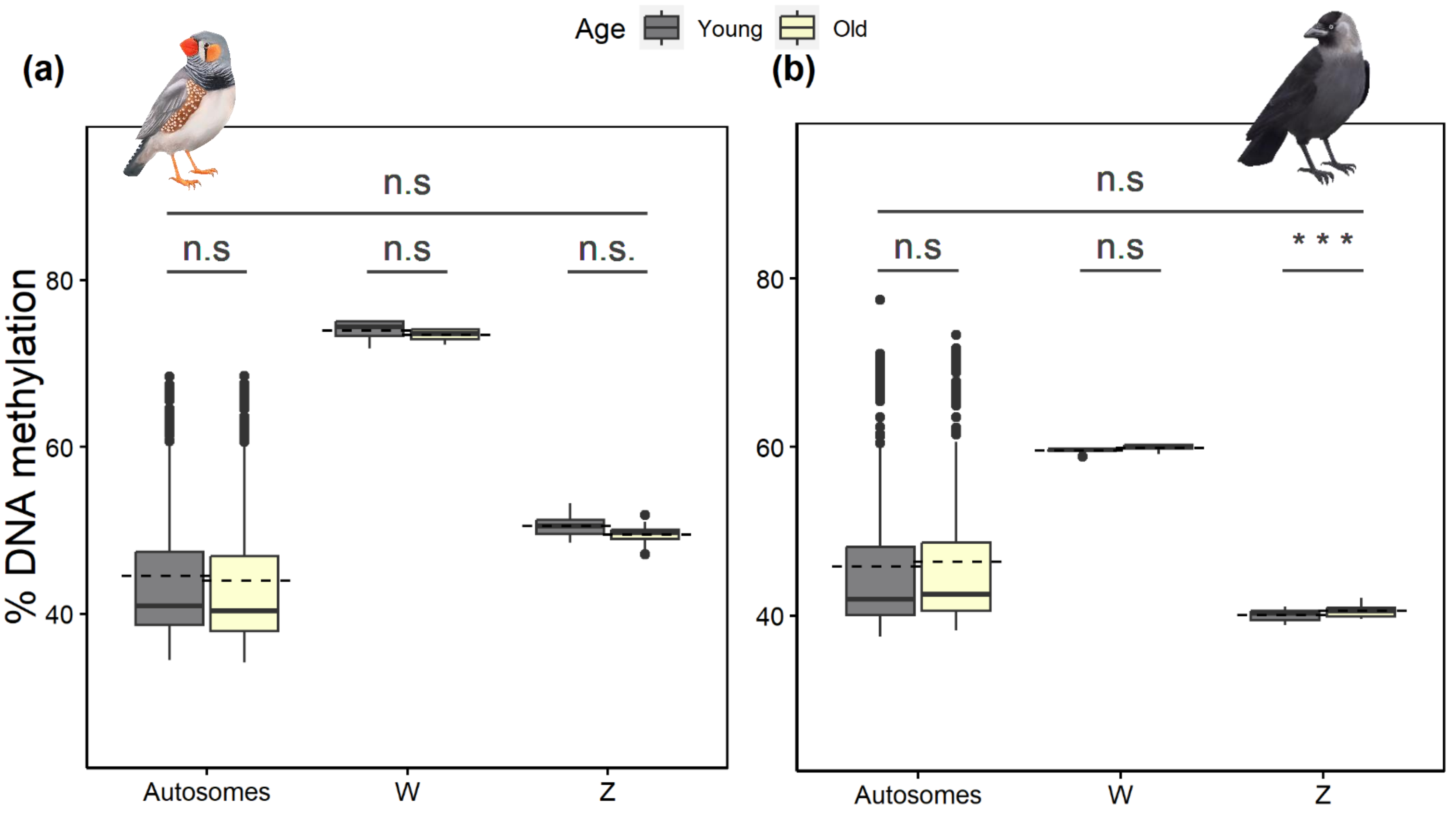

Fig. S1. Chromosome type-specific DNA methylation in old and young (a) zebra finches and (b) jackdaws. Boxplots on raw data represent median values, interquartile ranges and outliers (dots) and dashed lines represent the means. Data shown are based on the average DNA methylation per chromosome in each sample. Significance indicated by: *** $p < 0.001$, n.s. $p > 0.05$.

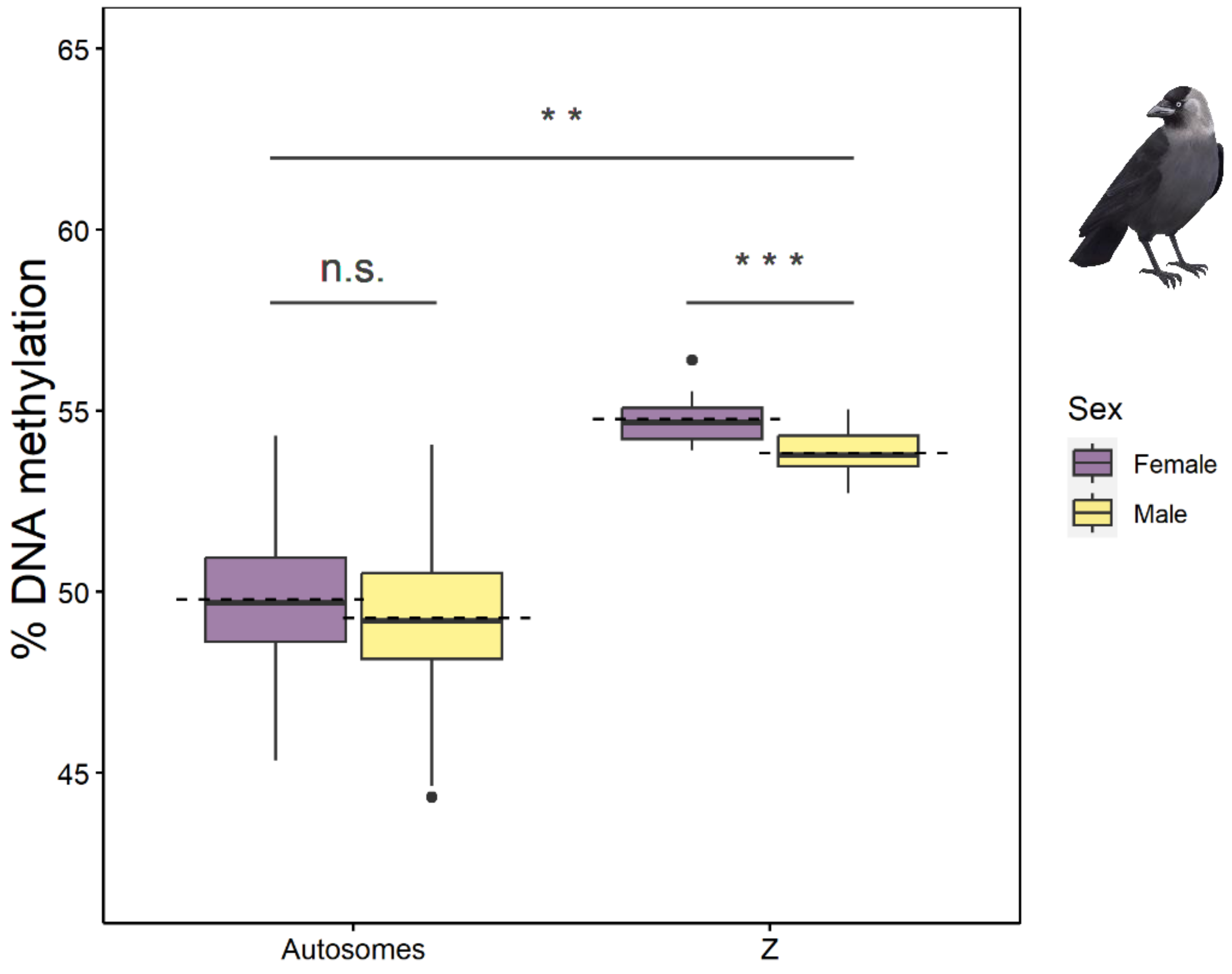

<u>Fig. S2. Chromosome type-specific DNA methylation in males and females in jackdaws aligned to jackdaw genome</u>. Boxplots on raw data represent median values, interquartile ranges and outliers (dots) and dashed lines represent the means. Data shown are based on the average DNA methylation per chromosome in each sample. Significance indicated by: *** $p < 0.001$, ** $p < 0.01$, n.s. $p > 0.05$.

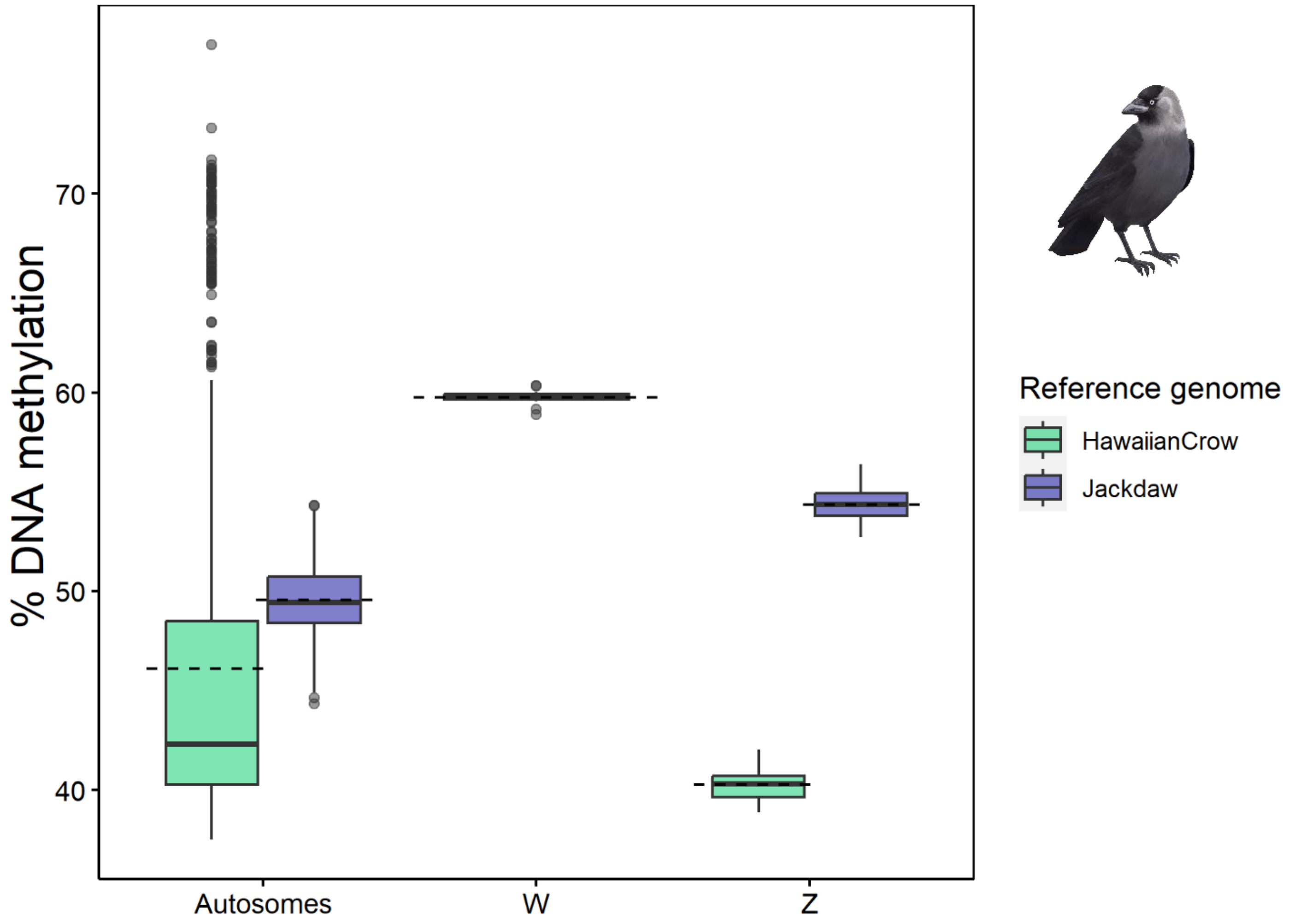

<u>Fig. S3. Chromosome type-specific DNA methylation in jackdaws aligned to Hawaiian crow and jackdaw genomes</u>. Boxplots on raw data represent median values, interquartile ranges and outliers (dots) and dashed lines represent the mean. Data shown are based on the average DNA methylation per chromosome in each sample.

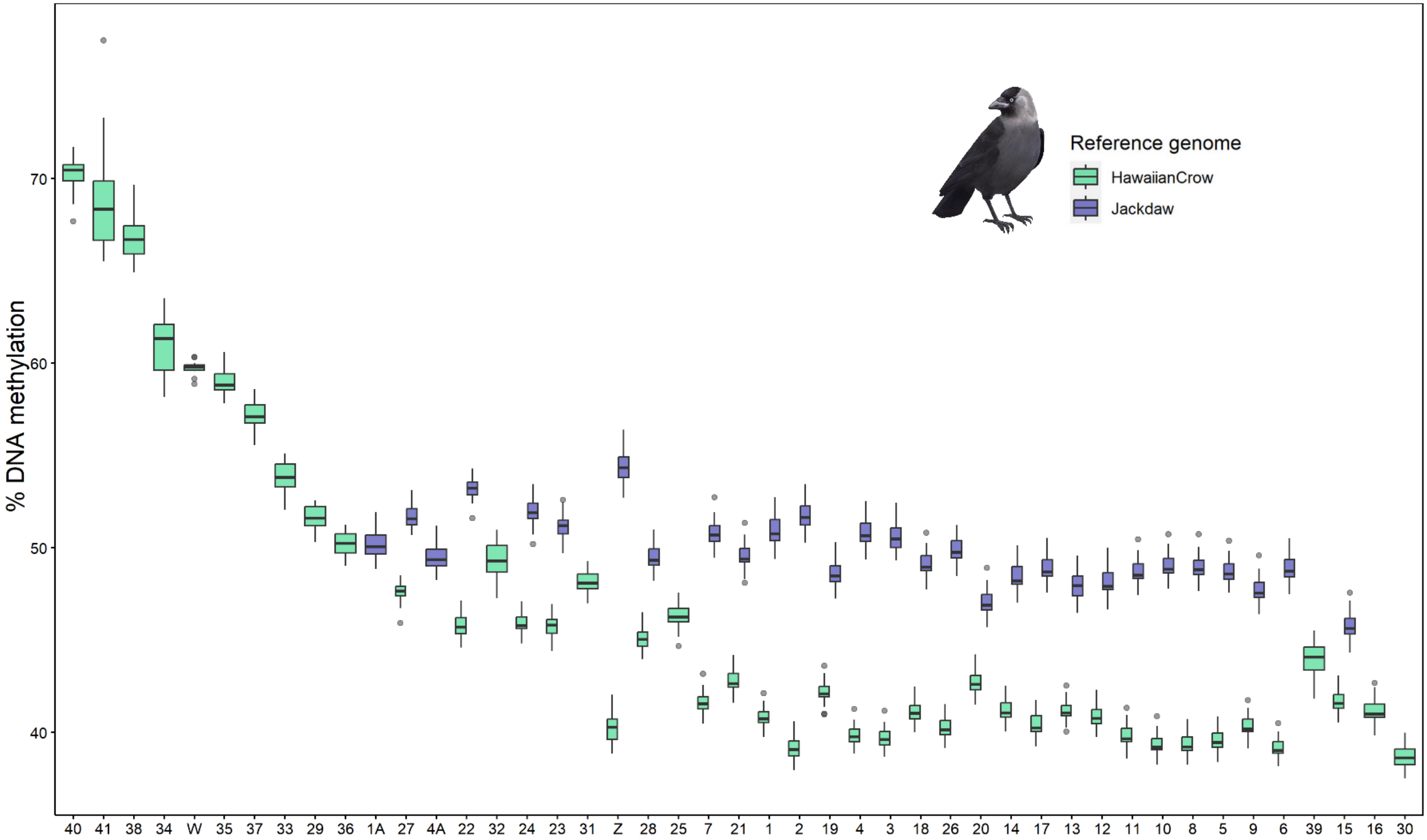

Fig. S4. Chromosome-wise DNA methylation levels in jackdaws aligned to Hawaiian crow and jackdaw genomes, ordered from the highest to lowest median value per chromosome along x axis. Boxplots on raw data represent median values, interquartile ranges and outliers (dots). Data shown are based on the average DNA methylation level per chromosome in each sample.

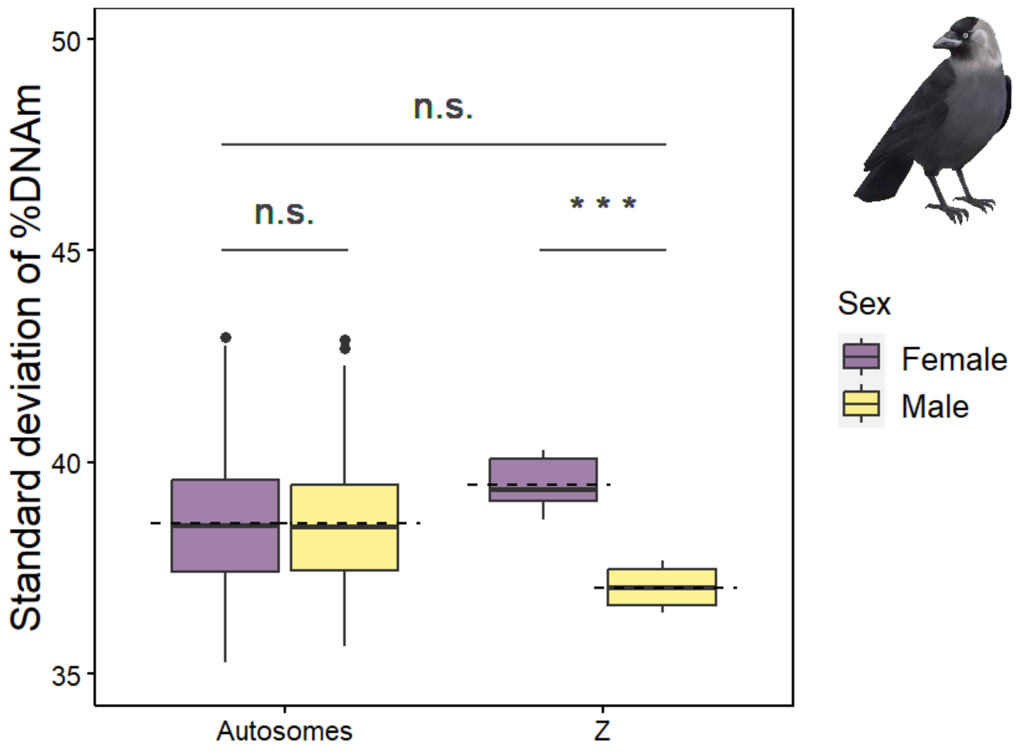

Fig. S5. Standard deviation of percentage of DNA methylation per chromosome in jackdaws aligned to jackdaw genome depending on the chromosome type and sex. Boxplots on raw data represent median values, interquartile ranges and outliers (dots) and dashed lines represent the mean. Significance indicated by: *** $p < 0.001$, n.s. $p > 0.05$.

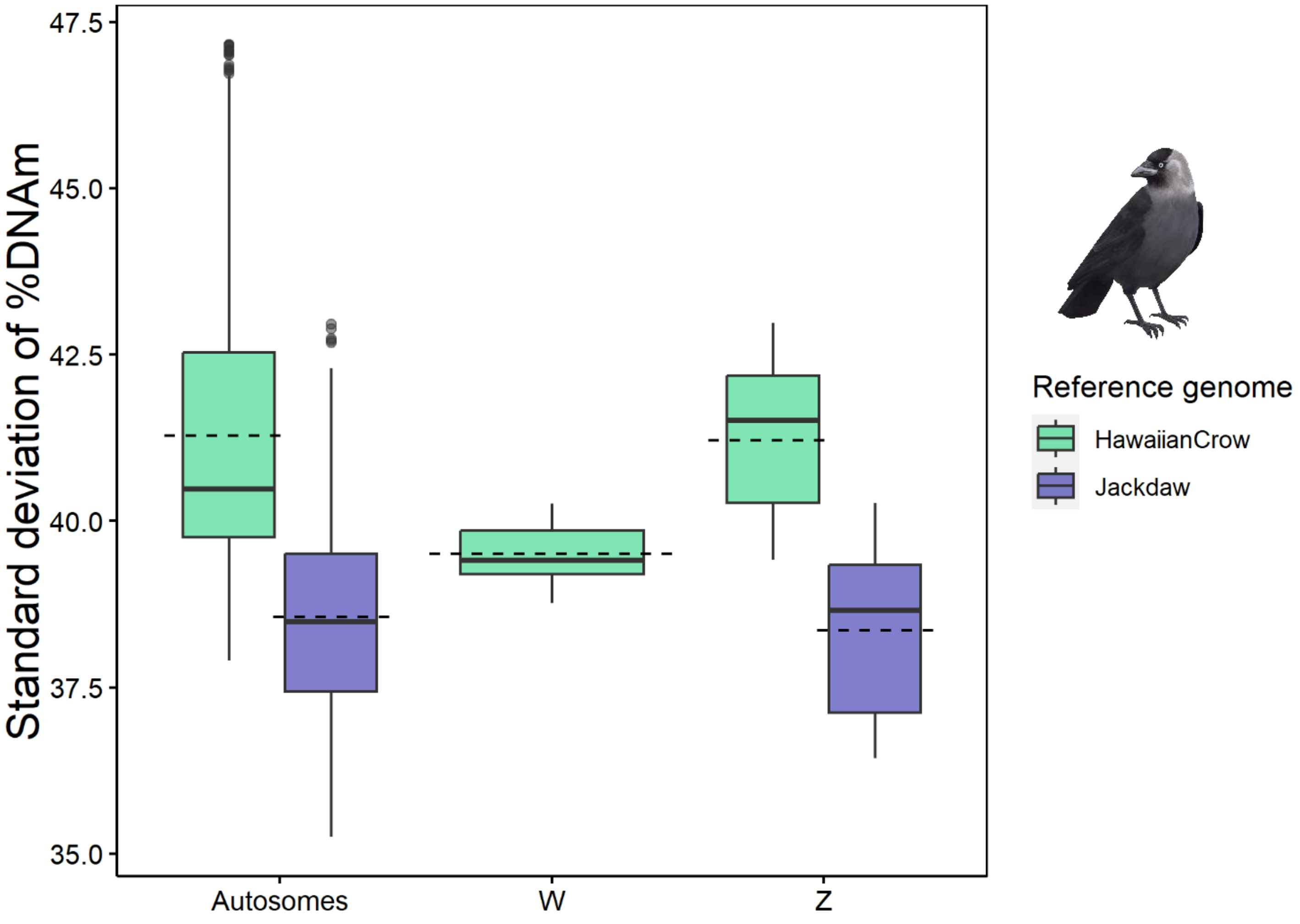

Fig. S6. Standard deviation of percentage of DNA methylation per chromosome in jackdaws aligned to Hawaiian crow and jackdaw genomes depending on the chromosome type. Boxplots on raw data represent median values, interquartile ranges and outliers (dots) and dashed lines represent the mean.

**(a)**

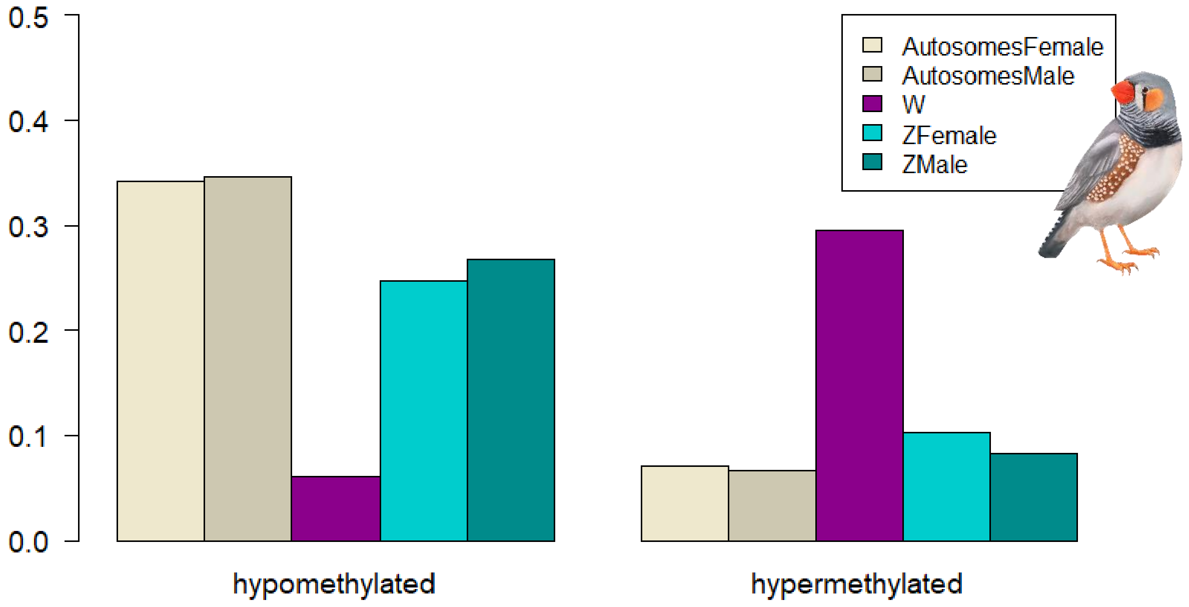

**(b)**

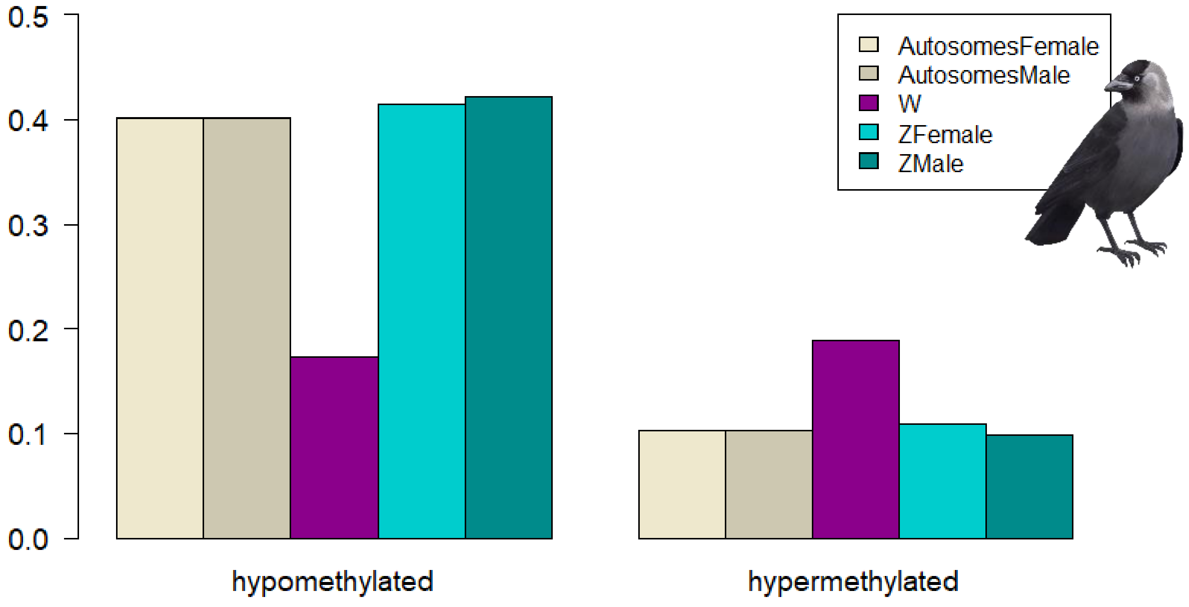

Fig. S7. Proportion of hypo- and hypermethylated sites to all sites per chromosome in zebra finches (a) and (b) jackdaws aligned to Hawaiian crow genome. DNA methylation level was averaged per each site across all samples that had this site available (weighted by coverage) to obtain hypomethylated (<=10% on average) and hypermethylated (>=90% on average) sites.
